# Targeting glutamine metabolism and redox state for leukemia therapy

**DOI:** 10.1101/303529

**Authors:** Mark A. Gregory, Travis Nemkov, Vadym Zaberezhnyy, Hae J. Park, Sarah Gehrke, Kirk C. Hansen, Angelo D’Alessandro, James DeGregori

## Abstract

Acute myeloid leukemia (AML) is a hematological malignancy characterized by the accumulation of immature myeloid precursor cells. AML is poorly responsive to conventional genotoxic chemotherapy and a diagnosis of AML is usually fatal. More effective and less toxic forms of therapy are desperately needed. AML cells are known to be highly dependent on the amino acid glutamine for their survival. Here, we show that blocking glutamine metabolism through the use of a glutaminase inhibitor (CB-839) significantly impairs antioxidant glutathione production in multiple types of AML, resulting in accretion of mitochondrial reactive oxygen species (mitoROS) and apoptotic cell death. Moreover, glutaminase inhibition makes AML cells susceptible to adjuvant drugs that further perturb mitochondrial redox state, such as arsenic trioxide (ATO) and homoharringtonine (HHT). Indeed, the combination of ATO or HHT with CB-839 exacerbates mitoROS and apoptosis, and leads to more complete cell death in AML cell lines, primary AML patient samples and *in vivo* using mouse models of AML. In addition, these redox-targeted combination therapies are effective in eradicating acute lymphoblastic leukemia cells *in vitro* and *in vivo*. Thus, targeting glutamine metabolism in combination with drugs that perturb mitochondrial redox state represents an effective and potentially widely applicable therapeutic strategy for treating multiple types of leukemia.

**Key Points:**

- Glutaminase inhibition commonly impairs glutathione metabolism and induces mitochondrial oxidative stress in acute myeloid leukemia cells
- A glutaminase inhibitor synergizes with pro-oxidant drugs in inducing apoptosis and eliminating leukemia cells *in vitro* and *in vivo*

## Introduction

Acute myeloid leukemia (AML) is a genetically heterogeneous disease that is characterized by the aberrant growth of myeloid progenitor cells with a block in cellular differentiation. AML is the most common adult acute leukemia and comprises 20% of childhood leukemias ^1,2^. AML is an aggressive malignancy and long-term (5 year) survival is less than 50% even for younger patients that can withstand intensive cytotoxic chemotherapy. Outcomes are far worse in elderly and less fit younger patients who can only receive less intensive chemotherapy (median survival < 1 year) ^3,4^. Significant improvement of AML patient outcomes will require the development of new therapeutic strategies that are both more effective and less toxic than currently used regimens. While in recent years there have been significant advances in our understanding of AML biology and the various genetic alterations that drive the disease, this knowledge has not, as of yet, led to development of targeted therapies that are capable of achieving long-term remissions. Moreover, for targeted therapies that have shown some modicum of clinical efficacy, such as those targeting FLT3 and IDH2, their utility is limited to subsets of AML that exhibit mutations in these genes^1,5,6^.

It has long been known that tumor cells rewire their metabolism to enable and sustain cell growth and proliferation and reprogrammed cellular metabolism is now recognized as a hallmark of cancer ^7–9^. It is feasible that the unique metabolic dependencies of tumor cells may be targeted for therapeutic effect. Many types of tumor cells, including AML, are highly dependent on glutamine for glutaminolysis, a mitochondrial pathway that involves the initial deamination of glutamine by the enzyme glutaminase *(GLS* and *GLS2*), yielding glutamate, which can then be converted into α-ketoglutarate to fuel the TCA cycle or used as a precursor for synthesis of glutathione, an important antioxidant factor ^10–13^ Our previous work has shown that inhibition of FLT3 in FLT3-mutated AML shuts down glutamine metabolism, leading to depletion of glutathione and cell death by apoptosis due to overwhelming oxidative stress ^14^ In addition, we and others have shown that inhibition of glutaminase in FLT3-mutated AML has similar effects as FLT3 inhibition on glutathione levels, redox state, and cell viability ^15,16^.

In the present study, we sought to determine if glutaminase inhibition could universally compromise glutathione metabolism and redox state in multiple types of AML and thus demonstrate broad therapeutic potential. Indeed, we found that a glutaminase inhibitor (CB-839) inhibits glutathione production, induces mitochondrial reactive oxygen species (mitoROS) and causes apoptotic cell death in many types of AML, as well as in acute lymphoblastic leukemia (ALL). Moreover, we demonstrate that combining a glutaminase inhibitor with a drug that further induces mitoROS is an effective therapeutic strategy for eliminating leukemia cells both *in vitro* and *in vivo*.

## Materials and Methods

### Cell culture

The AML cell lines Molm13 and MV4-11 (FLT3-ITD^+^), EOL-1 (FLT3^WT^ but FLT3-dependent ^17^), HL-60, Kasumi-1, OCI-AML3, THP-1 and U937 were obtained from D. Graham (Emory University SOM). MonoMac6 (FLT3^V592A^) was obtained from K. Bernt (University of Colorado SOM). KG1a was obtained from Gail Eckhardt (University of Texas at Austin Dell SOM). The CML cell line K562 and ALL cell line SUP-B15 were purchased from ATCC. Z-119 Ph^+^ ALL cells were obtained from Z. Estrov (MD Anderson). All cell lines were cultured in RPMI 1640 medium/10% FBS, except for HL-60 (Iscove’s Modified Dulbecco’s Medium [IMDM]/20% FBS), K562 (IMDM/10% FBS) and SUP-B15 and Z-119 (RPMI 1640/20% FBS with 5mM β-mercaptoethanol). Cell lines were authenticated by short tandem repeat examination and tested negative for mycoplasma using the e-Myco plus PCR Detection Kit (iNtRON). Murine Bcr-Abl^+^ ARF-/- ALL cells ^18^ were cultured in RPMI 1640 medium/10% FBS and 2x L-glutamine and murine MLL-ENL/FLT3-ITD^+^ AML cells ^19^ in 50% IMDM/RPMI 1640 with 20% FBS prior to inoculation into mice.

### Pharmacologic agents

CB-839 was purchased from MedChem Express and, for in vivo experiments, was obtained from Calithera Biosciences (South San Francisco, CA).

Homoharringtonine (HHT; omacetaxine mepesuccinate), arsenic trioxide (ATO), cell-permeable glutathione reduced ethyl ester (GSH-MEE) and dimethyl 2-oxoglutarate (α-ketoglutarate) were purchased from Millipore Sigma.

### Metabolic tracing experiments

Cells were seeded at 3 × 10^5^/ml (replicates of 3) and treated with vehicle (DMSO) or CB-839 at 500 nM for 8 h, followed by incubation in glutamine-free RPMI 1640 supplemented with ^13^C_5_,^15^N_2_-labeled L-glutamine (Cambridge Isotope Laboratories) for up to 12 h, in the presence of vehicle or drug. Flash frozen cell pellets (~ 1 × 10^6^ cells) or supernatants (50 μl) were extracted and subjected to analysis by ultra-high pressure liquid chromatography and mass spectrometry (UHPLC/MS) as was previously described ^20^. Metabolite assignments, isotopologue distributions, and correction for expected natural abundances of ^13^C and ^15^N isotopes were performed using MAVEN (Princeton University, Princeton, NJ) and manually validated.

### Cell viability assays

Cells were seeded at 0.5-1.0 × 10^5^/ml in triplicate wells of 48-well tissue culture plates. Where indicated, the cells were treated with drug for a period of 48-72 h. After treatment, a sample of cells from each well was stained with PI (10 μg/ml) and viable cells (PI^−^) were counted with a flow cytometer (Millipore Guava easyCyte 8HT). Alternatively, cells were stained using 7-AAD/anti-Annexin V (Nexin reagent, Millipore Sigma) to detect apoptotic cells. ATP levels were measured using the Cell Titer-Glo Assay (Promega).

### Viability assays on primary AML samples

Primary AML patient samples were obtained from C.T. Jordan (University of Colorado SOM). Total bone marrow mononuclear cells were isolated by standard Ficoll procedures (GE Healthcare) and cryopreserved in freezing medium consisting of Cryostor CS10 (BioLife Solutions). The viability of leukemic cells after thawing was >90%. The cells were cultured in IMDM media with 4mM glutamine and 10% heat-inactivated FBS and supplemented with 10 μg/ml of IL-3, IL-6, IL-7, GM-CSF, and SCF (Peprotech) for 1 day prior to exposure to drugs. After 72 h of drug treatment, viable cells (PI-excluded) were counted by flow cytometry.

### ROS and glutathione measurements

Cells (1 × 10^5^) were treated with drug as indicated, washed in PBS, resuspended in 150 μL of 5 μM 2’,7’-dichlorodihydrofluorescein diacetate (H_2_DCFDA; Thermo Fisher Scientific) to measure global peroxide levels, or 10 μM MitoPY1 (Tocris Bioscience) to measure mitochondrial peroxide (MitoROS) levels, in serum-free media, and incubated at 37°C for 15 or 60 min, respectively.

Cells were washed in cold FACS buffer, re-suspended in FACS buffer and immediately analyzed by flow cytometry. To measure glutathione levels, the GSH-Glo Assay (Promega) was utilized according to the manufacturer’s instructions.

### Mouse experiments

NOD.Cg-*Prkdc^scid^ ll2rg^tm1Wjl^/SzJ* (NSG) were obtained from The Jackson Laboratory and bred in house. The patient sample for xenograft (obtained from D. Pollyea, University of Colorado SOM) came from a 54-year old female with AML expressing FLT3-ITD and NPM1 mutations. The leukemia was engrafted into female NSG mice as previously described ^14^ Treatment was started when peripheral blast count was 3-10% (mean of 6% for all groups). For monitoring leukemic burden in mice, blood was stained with anti-human CD45-FITC and HLA-ABC-PE-Cy7 (BD Biosciences) and analyzed by flow cytometry. For the MLL-ENL/FLT3-ITD AML and Bcr-Abl^+^ ALL models, 6-10 week old female C57BL/6 mice (National Cancer Institute) were utilized. C57BL/6 mice were inoculated intravenously with 1 × 10^6^ MLL-ENL/FLT3-ITD AML luciferase cells or 5 × 10^5^ ARF-/- p185 Bcr-Abl/GFP B-ALL cells. After 3-5 days, treatment was started. Treatment at the indicated doses of drugs was 5 days on/2 days off. In the ALL model, leukemic burden was monitored by quantifying GFP/B220^+^ and Mac1^−^ blasts in the peripheral blood as previously described^21^. In the AML model, leukemic burden was monitored by bioluminescent imaging using a Xenogen IVIS200 imaging device and standard protocols. For in vivo delivery, CB-839 was solubilized in in 25% HP-β-CD/10 mM citrate pH 2.0 (vehicle), and ATO and HHT in sterile saline. The Institutional Animal Care and Use Committee at University of Colorado approved all mouse experiments under protocol #41414(05)1 E.

### Statistics

All data are expressed as the mean ± SEM. Comparisons between two values were performed by Mann Whitney *U* test, significance was defined at *P* ≤ 0.05 *, *P* ≤ 0.01 **. Combination indices were calculated using the median-effect principle and the Combination Index-Isobologram Theorem (CompuSyn).

## Results

Our previous studies have shown that impeding glutamine uptake or metabolism using either a FLT3 inhibitor or glutaminase inhibitor, respectively, impairs anti-oxidant defenses by limiting glutathione synthesis and leads to the accumulation of mitochondrial oxidative stress and subsequent apoptosis in FLT3-mutated AML cells ^14,15^. FLT3, however, is mutated in only a subset of AMLs. Thus, we wanted to determine if glutaminase inhibition could similarly impair glutamine/glutathione metabolism in AML cells of other genotypes, indicating the potential for broader therapeutic utility. To this end, we employed metabolic flux analysis using ^13^C, ^15^N-labeled glutamine (Figure S1A) in two FLT3 wild type (WT) AML cell lines, OCI-AML3 and EOL-1. These cells were treated with vehicle or the glutaminase inhibitor CB-839 for 8 h hollowed followed by incubation with labeled glutamine (up to 12 h). As shown in Figure 1A, the cells treated with CB-839 displayed severe impairment of glutamine flux into glutamate and its downstream metabolites aspartate, alanine and α-ketoglutarate. Since α-ketoglutarate feeds into the tricarboxylic (TCA) cycle, flux into TCA cycle intermediates (succinate, malate, and citrate) was severely reduced. As was previously observed in FLT3 mutated AML, CB-839 also impaired flux into antioxidant glutathione, leading to lower overall glutathione levels (Figure S1B).

**Figure 1.**
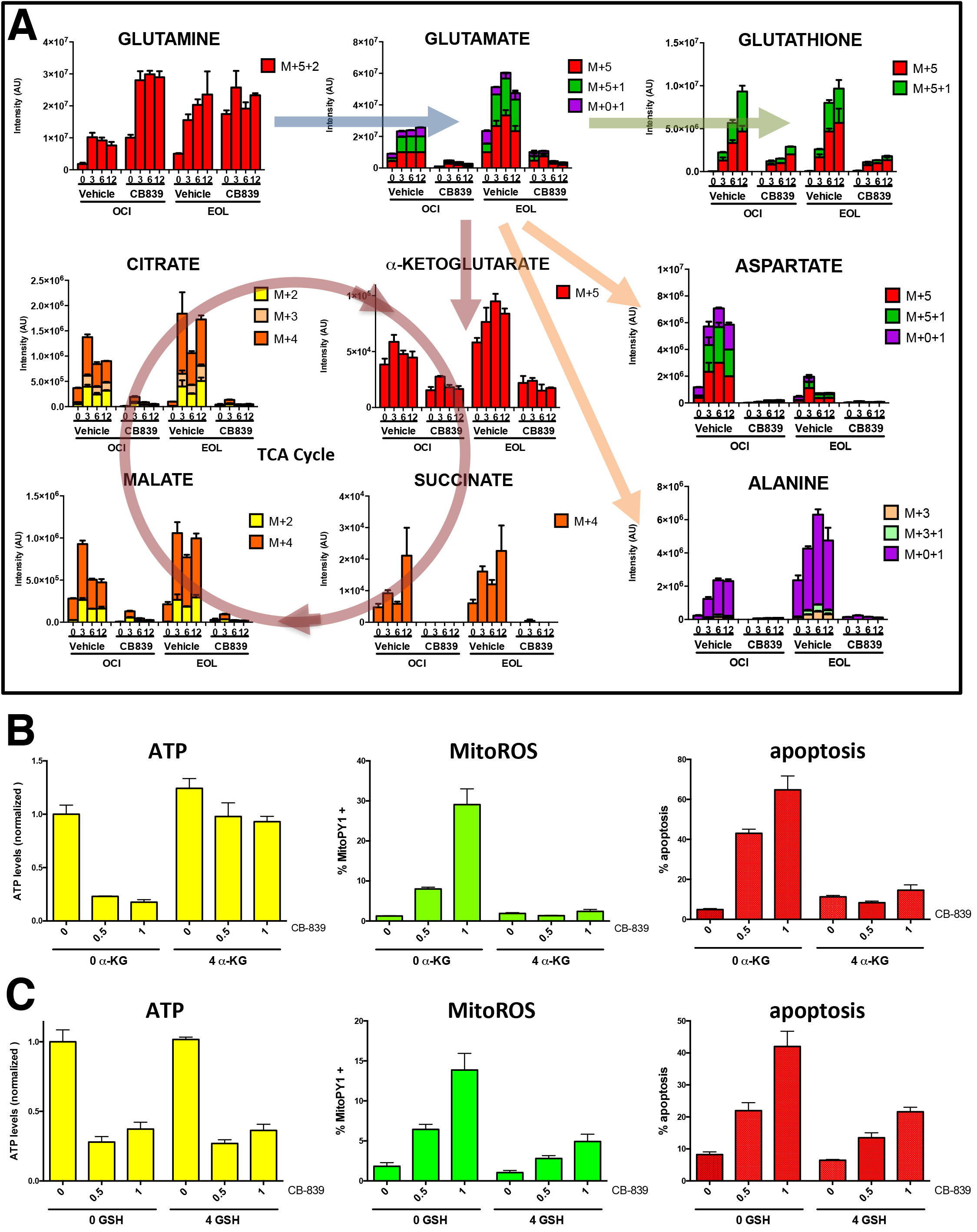
The glutaminase inhibitor CB-839 impairs energy and redox metabolism in FLT3^WT^ AML cells. **A)** Glutamine flux analysis of OCI-AML3 and EOL-1 AML cells treated with vehicle (DMSO) or CB-839 (500 nM). Cells were pre-treated for 8 h followed by ^13^C,^15^N-glutamine labeling for 0, 3, 6, and 12 h, as indicated, followed by UHPLC/MS analysis. Levels of ^13^C and^15^N containing isotopologues (based on signal intensity) are shown. **B,C)** EOL-1 cells were treated with CB-839 (0, 0.5, or 1 μM, as indicated) with simultaneous addition of cell-permeable α-ketoglutarate (4 mM α-KG; **B**) or cell-permeable glutathione (4 mM GSH; **C**) for 24 h and ATP levels and MitoROS levels (using the dye MitoPY1) were measured. Apoptosis was measured after 48 h.

The depletion of α-ketoglutarate and glutathione could both contribute to CB-839-induced apoptosis, through energy (ATP) depletion due to TCA cycle interference and/or through oxidative stress due to lack of anti-oxidant capacity. In attempt to parse the importance of the TCA cycle for cell survival, we added dimethyl-glutarate to EOL-1 cells upon treatment with CB-839. Dimethyl-glutarate is a membrane-permeable dimethyl-ester of α-ketoglutarate (α-KG), which is metabolized to α-KG by cellular esterases. As shown in Figure 1B, as expected, α-KG was capable of reversing ATP depletion caused by CB-839 and also substantially suppressed the induction of apoptosis. Unexpectedly, however, α-KG also substantially suppressed the induction of mitochondrial reactive oxygen species (mitoROS) caused by CB-839, indicating the possibility that exogenous α-KG could be used for generation and maintenance of glutathione. Indeed, α-KG was able to prevent depletion of glutathione in EOL-1 cells upon treatment with CB-839 (Figure S1C). To parse the importance of glutathione for cell survival, we added cell-permeable glutathione (GSH-MEE) to EOL-1 cells upon treatment with CB-839. GSH-MEE was completely incapable of rescuing ATP levels (Figure 1C) and while treatment with exogenous GSH-MEE was only capable of partial maintenance of glutathione levels (Figure S1C), this was sufficient to significantly suppress induction of MitoROS and subsequent apoptosis (Figure 1C). These results suggest a prominent role for glutathione depletion and the induction of mitoROS in the causation of apoptosis and AML cell death upon glutaminase inhibition.

To determine if glutathione depletion is a universal consequence of glutaminase inhibition in AML, we treated a panel of AML cell lines (EOL-1, Molm13, MonoMac6, MV4-11, HL-60, Kasumi-1, OCI-AML3, and THP-1), along with U937 (lymphoma) and K562 (chronic myeloid leukemia; CML) cell lines with CB-839 for 24 h. As shown in Figure 2A, CB-839 caused a reduction in glutathione levels (normalized to untreated cells; unnormalized data shown in Figure S2A) in all AML cell lines examined while this was not observed in U937 and K562 cells. Moreover, the degree of glutathione depletion almost perfectly mirrored the amount of mitoROS (Figure 2B) and apoptosis (Figure 2C) induction caused by CB-839 treatment. On the contrary, there was no correlation with overall cellular levels of reactive oxygen species (Figure S2B). These results again show that maintenance of mitochondrial redox state is critical for AML cell survival and suggest a novel therapeutic strategy (Figure 2D): A glutaminase inhibitor, while able to induce some degree of mitoROS and apoptosis by itself through inhibition of glutathione production, may sensitize AML cells to a second mitoROS-inducing drug. The combination of these drugs would result in the superinduction of mitoROS leading a more complete killing of the leukemia cells.

**Figure 2.**
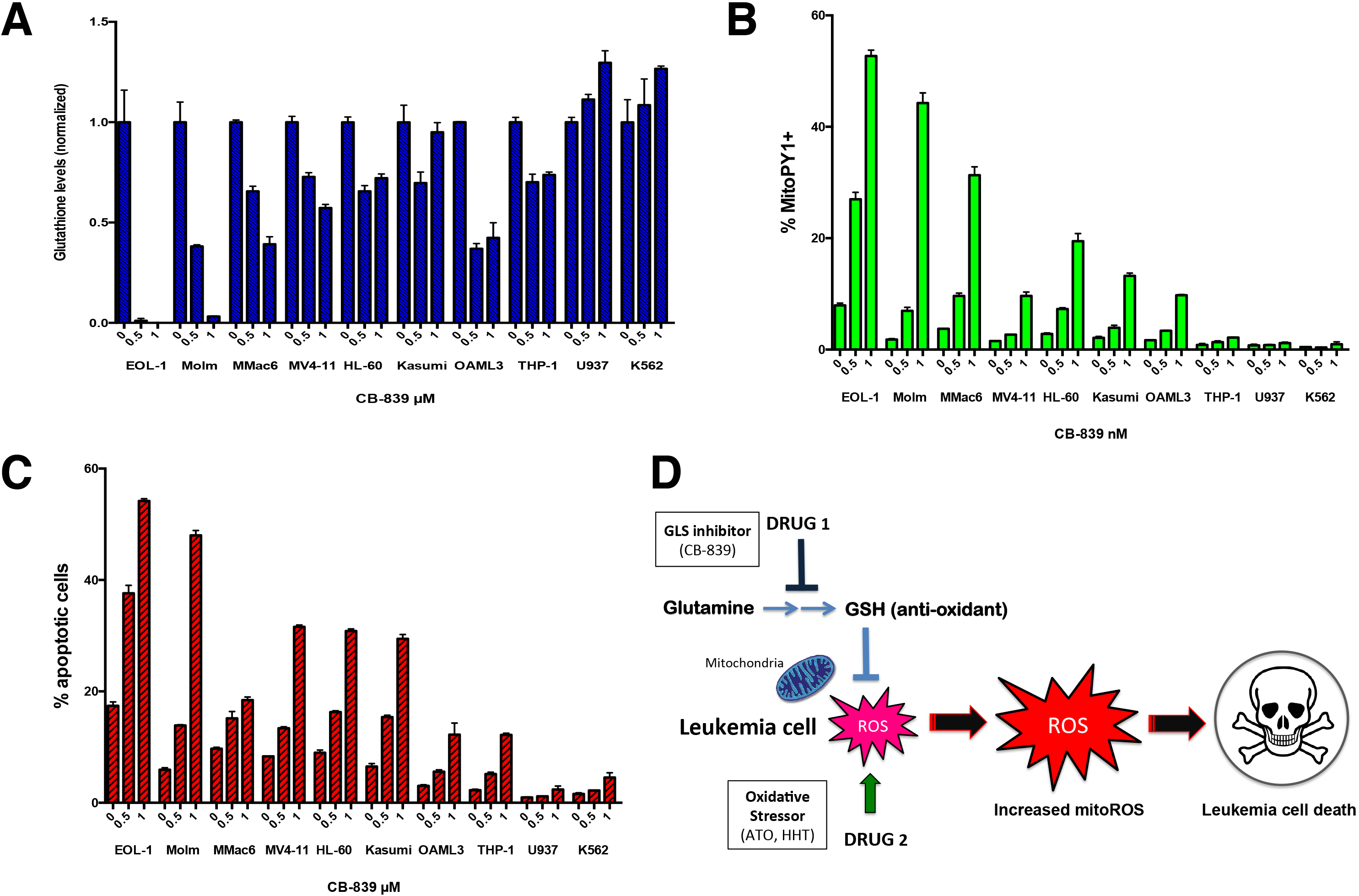
The glutaminase inhibitor CB-839 decreases glutathione levels, and induces mitoROS and apoptosis in AML cells. **A,B,C)** The AML cell lines EOL-1, Molm13, MonoMac6, MV4-11, HL-60, Kasumi-1, OCI-AML3, and THP-1, and the lymphoma-derived cell line U937 and the CML cell line K562 were treated with CB-839 (0, 0.5, or 1 μM, as indicated) and glutathione (**A**) and mitoROS (**B**) levels (using the dye MytoPY1) were measured after 24 h. Apoptosis was measured after 48 h (**C**). **D)** Model showing mechanism of redox-directed combination therapy for treating leukemia. Use of a glutaminase inhibitor (Drug 1, e.g. CB-839) blocks glutamine metabolism and production of anti-oxidant GSH and makes leukemia cells vulnerable to a pro-oxidant agent (Drug 2, e.g. ATO or HHT), leading to superinduction of mitoROS and subsequent leukemia cell death via apoptosis.

In order to test the therapeutic strategy described above, we first tested the efficacy of arsenic trioxide (ATO) combined with CB-839 on AML cell lines *in vitro*. ATO is an agent that promotes generation of ROS and mitochondrial apoptosis, and it has long been known to have anti-leukemic activity^22^. As shown in Figure 3A, ATO by itself induces mitoROS in a dose-dependent manner; however, ATO induces higher levels of mitoROS when used in combination with CB-839. Moreover, ATO and CB-839 synergized in killing EOL-1 cells (Figure 3B). Combination index (CI) values were synergistic (< 0.6) at all combination doses tested and were strongly synergistic (< 0.3) with most combinations (all CI values listed in Table S1). Effects on viability correlated with increased apoptosis (Figure 3C). Comparable results were obtained using Molm13 cells; while ATO did not induce appreciable mitoROS in these cells by itself, it strongly induced mitoROS when used in combination with CB-839 (Figure 3D), which correlated with increased cell killing (Figure 3E) and apoptosis (Figure 3F). This drug combination had similar effects on cell viability and apoptosis in several additional AML cell lines, including OCI-AML3, MV4-11, KG1a, HL-60, and MonoMac6 (Figures S3A-F).

**Figure 3.**
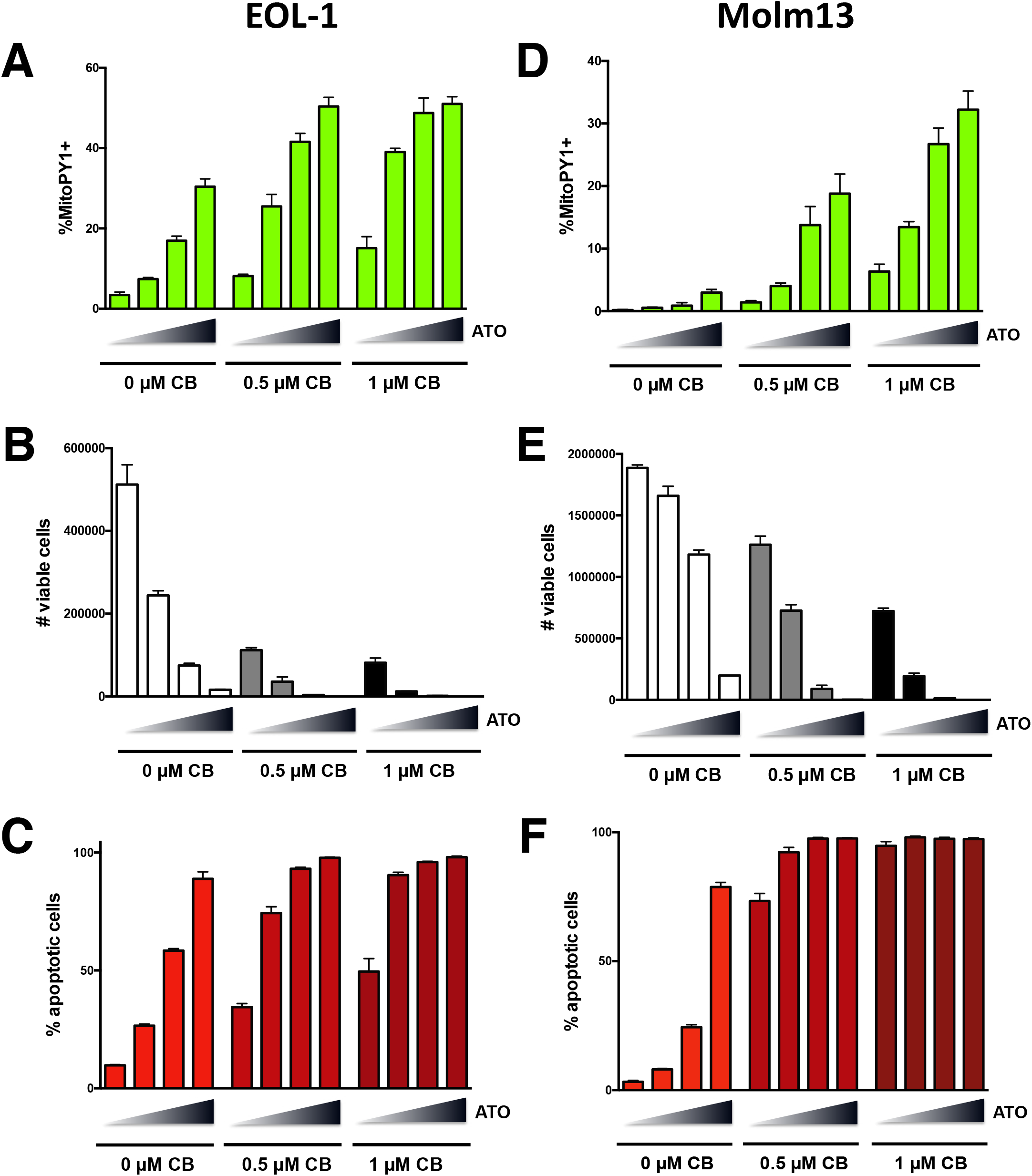
CB-839 cooperates with the pro-oxidant drug ATO in inducing mitoROS, apoptosis, and AML cell death. EOL-1 (**A,B,C**) or Molm13 (**D,E,F**) were treated with CB-839 and ATO alone or together as indicated (increasing concentrations of ATO of 0, 0.2, 0.4, and 0.8 μM indicated by triangles) and mitoROS levels were measured using MitoPY1 (**A,D**). After 72 h, the number of viable cells (based on PI exclusion) was counted by flow cytometry (**B,E**) and apoptosis was measured after 48 h (**C,F**).

We next wanted to test the therapeutic efficacy of CB-839 combined with ATO *in vivo*. For this purpose, we employed a genetically engineered mouse model of MLL-ENL/FLT3-ITD-initiated AML^19^. Briefly, fetal liver cells isolated from C57Bl/6 (B6) mice were transduced with retroviruses encoding MLL-ENL and FLT3-ITD together with luciferase. Transplantation of these cells generates a myeloid leukemia similar to human AML. These primary mouse AML cells, which can be cultured *in vitro*, are then capable of transferring disease to unconditioned B6 recipient mice with high penetrance. Three days after transferring the primary AML cells to recipient mice, therapy was initiated using vehicle, ATO, CB-839, or CB-839 and ATO in combination (n=5 for each group). Leukemic burden was quantified using bioluminescent imaging on days 4 and 11 after the start of therapy (Figure 4A). By day 11, leukemic burden was significantly lower in the combination treated mice compared to the groups treated with either ATO or CB-839 alone (Figure 4A and B), indicating superior therapeutic efficacy.

**Figure 4.**
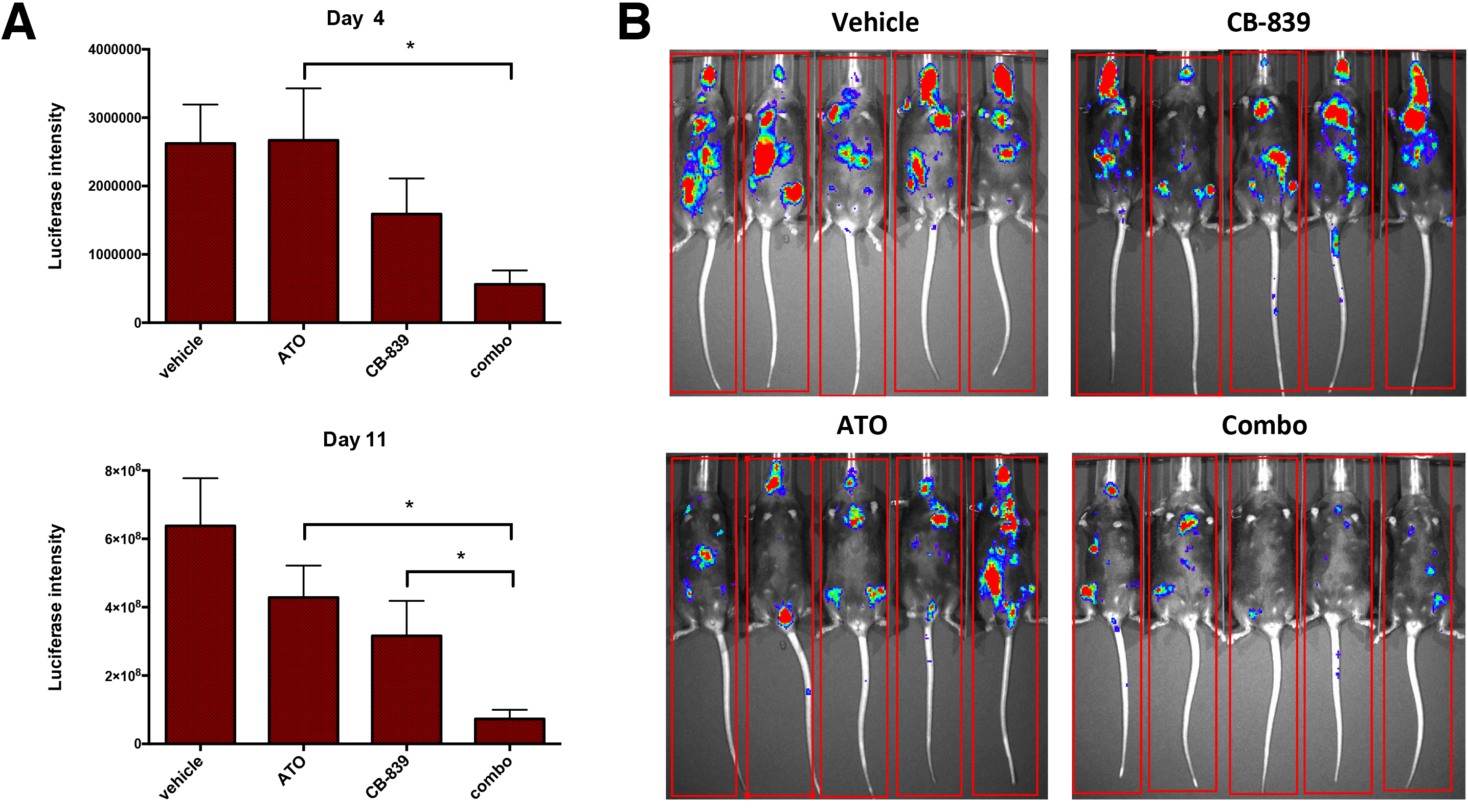
CB-839 cooperates with ATO in eliminating AML cells *in vivo*. MLL-ENL/FLT3-ITD^+^ luciferase expressing mouse leukemia cells were transplanted into Bl6 mice and after 3 days treatment was initiated with vehicle (n=5), CB-839 (n=5; 200 mg/kg twice daily p.o.), ATO (n=5; 8 mg/kg once daily i.p.), or CB-839 and ATO in combination (n=5) for up to 11 days (4 days on/2 days off/ 5 days on therapy). **A)** Leukemic burden was measured by bioluminescence using an in vivo imaging system (IVIS) on the indicated days. **B)** IVIS images from Day 11.

To determine if CB-839 will effectively combine with another drug that induces mitochondrial ROS, we next explored homoharringtonine (HHT). HHT is a protein translation inhibitor indicated for the treatment of tyrosine kinase inhibitor-resistant CML ^23^ that was recently found to effectively combine with FLT3 inhibitors for the treatment of AML ^24^. We have previously shown that drugs that combine well with FLT3 inhibitors are frequently potent inducers of mitoROS ^14^. Indeed, we found that HHT strongly induced mitoROS in Molm13 AML cells and mitoROS was exacerbated when HHT was used in combination with CB-839 (Figure 5A). In addition, HHT synergized with CB-839 in killing Molm13 cells (Figure 5B), which correlated with induction of apoptosis (Figure 5C). Similar effects on cell viability using this drug combination were observed in EOL-1 cells (Figure 5D). To test efficacy *in vivo*, we utilized the MLL-ENL/FLT3-ITD AML mouse model described above. Five days after leukemia transfer, therapy was initiated using vehicle, CB-839, HHT, or CB-839 and HHT in combination (n=5 per group). As shown in Figure 5E, by Day 8 of therapy, leukemic burden was significantly lower in the combination group compared to the CB-839 and HHT groups, and remained significantly lower on Day 15 (Figure 5E and F). These results demonstrate the therapeutic efficacy of the CB-839 plus HHT combination *in vivo*, at least in this model of AML.

**Figure 5.**
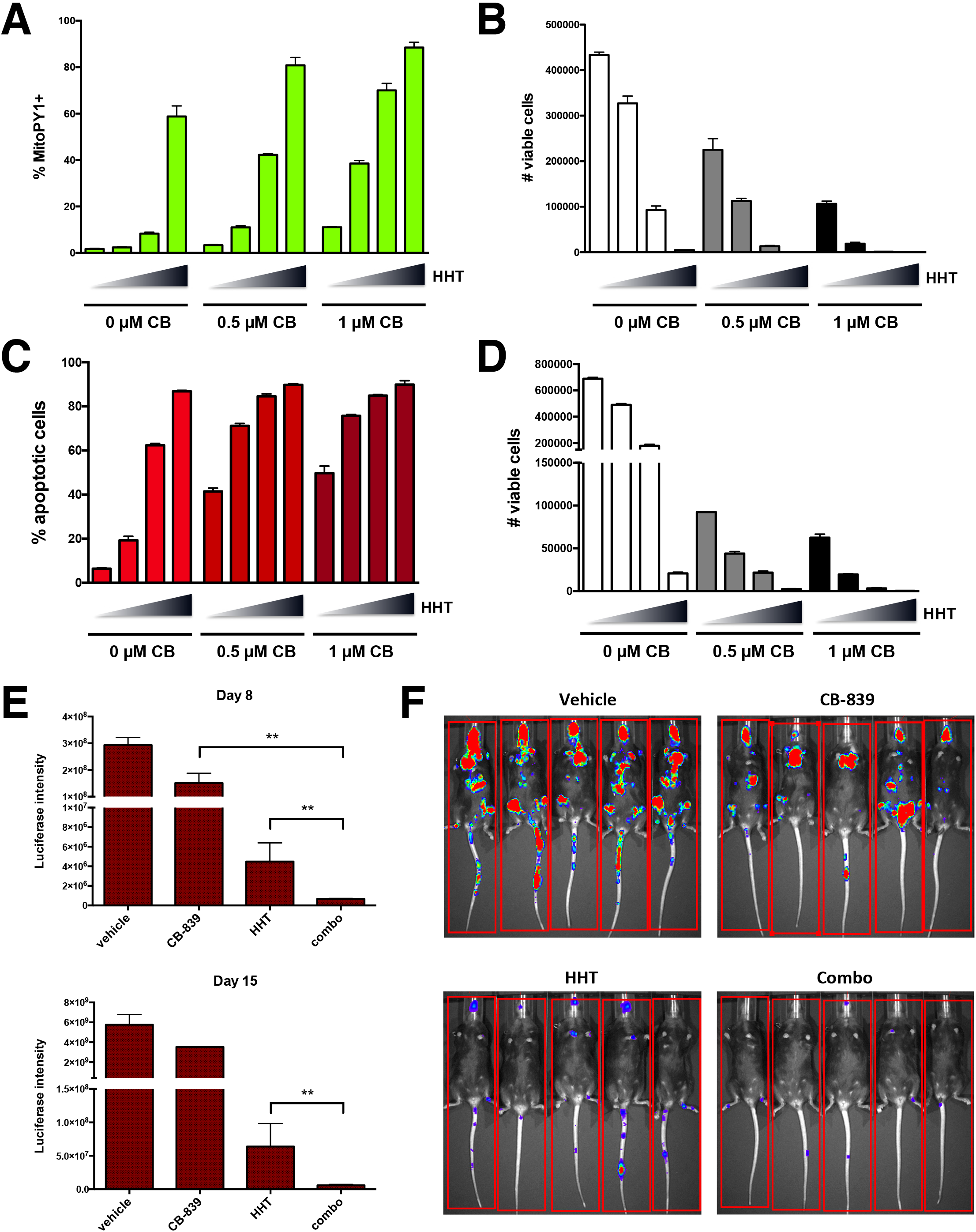
HHT induces mitoROS and cooperates with CB-839 in eliminating AML cells *in* vitro and *in vivo*. **A,B,C)** Molm13 cells were treated with CB-839 and HHT alone or together as indicated for 20 h (triangles: 0, 5, 10, and 20 nM HHT) and mitoROS levels were measured using MitoPY1 (**A**). After 48h, the number of viable cells (based on PI exclusion) was counted by flow cytometry (**B**) and apoptosis was measured after 24 h (**C**). **D)** EOL-1 cells were treated with CB-839 and HHT alone or together as indicated for 72 h (triangles: 0, 5, 10, and 20 nM HHT) and the number of viable cells (based on PI exclusion) was counted by flow cytometry. **E)** MLL-ENL/FLT3-ITD^+^ luciferase expressing mouse leukemia cells were transplanted into Bl6 mice and after 4 days, treatment was initiated with vehicle (n=5), CB-839 (n=5; 200 mg/kg twice daily p.o.), HHT (n=5; 1 mg/kg once daily i.p.), or CB-839 and HHT in combination (n=5) for up to 15 days (5 days on/2 days off therapy). Leukemic burden was measured by bioluminescence using an in vivo imaging system (IVIS) on the indicated days. **F)** IVIS images from Day 8.

To determine if CB-839 plus HHT shows anti-leukemic activity in human primary AMLs, we tested the effects of this combination on three AML patient samples, all of which are WT for FLT3. As shown in Figure 6, HHT synergized with CB-839 in AML cell killing. In addition, we show that the drugs cooperate in inducing mitoROS (Figure S4A) and apoptosis (Figure S4B), which corresponds with the observed decrease in cell viability (Figure S4C) in primary AML cells. To determine if CB-839 and HHT cooperate in eliminating primary AML samples *in vivo*, we utilized a patient-derived xenograft mouse model as previously described ^14^ Briefly, primary AML cells were engrafted into immunodeficient NSG mice, and after ~4 weeks, leukemic burden in the peripheral blood (PB) was determined by flow cytometry, the mice were divided into groups with approximately equal mean leukemic burden (~6%), and therapy was initiated using vehicle (n=5), CB-839 (n=5), HHT (n=5), or CB-839 and HHT in combination (n=6). Figure 6D shows that after 11 days of therapy, leukemic burden in PB was significantly lower in the combination group compared to the individual treatment groups and remained lower out to day 25, again demonstrating the superiority of the CB-839/HHT combination in treating AML.

**Figure 6.**
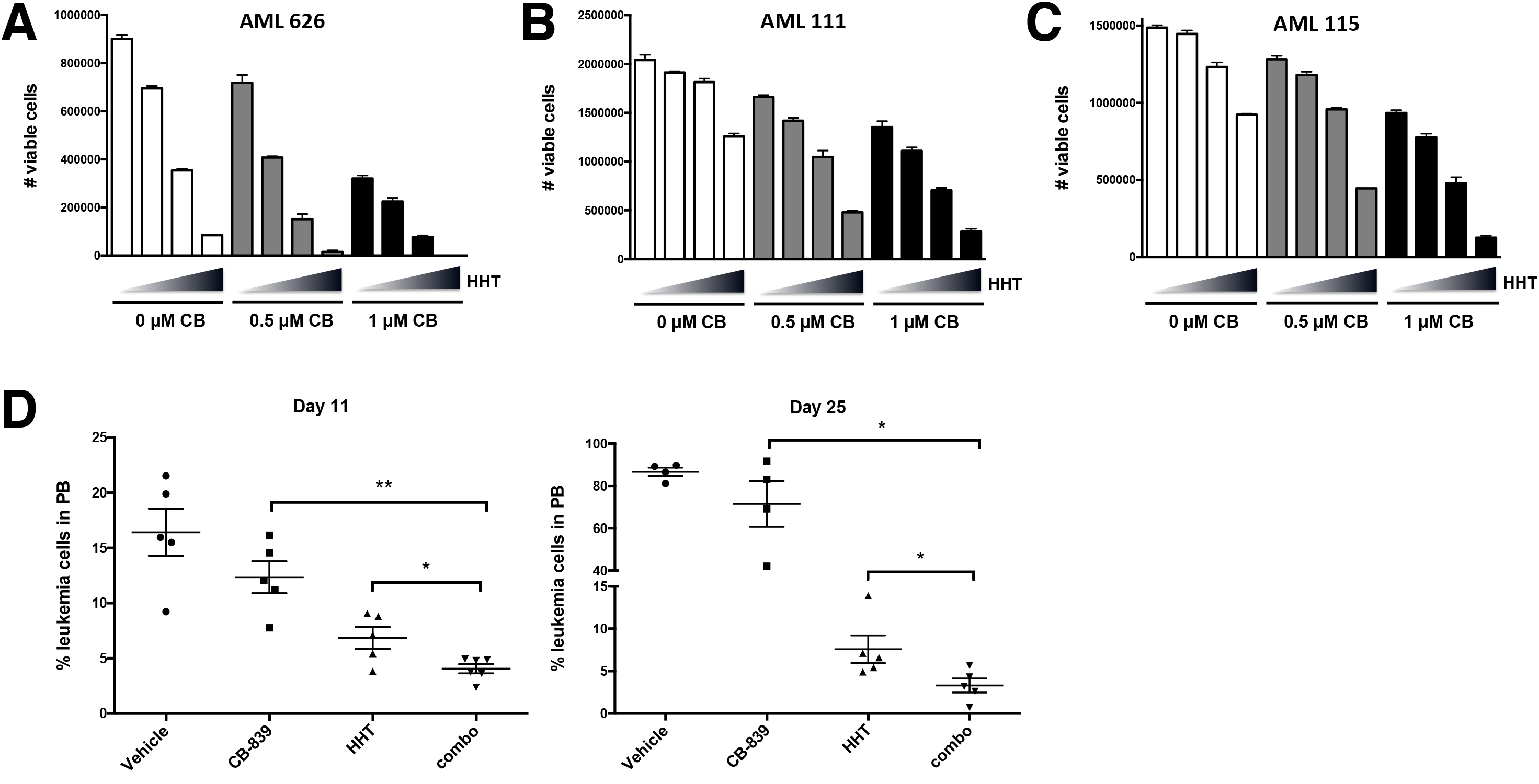
CB-839 cooperates with HHT in eliminating primary human AML cells *in* vitro and *in vivo*. **A,B,C)** Primary AML cells from three different patients (626, 111, 115) were treated with CB-839 and HHT alone or together as indicated for 72 h (triangles: 0, 5, 10, and 20 nM HHT) and the number of viable cells (based on PI exclusion) was counted by flow cytometry. **D)** NSG mice were engrafted with primary human AML cells and groups of mice were treated with vehicle (n=5), CB-839 (n=5; 200 mg/kg twice daily p.o.), HHT (n=5; 1 mg/kg once daily i.p.), or CB-839 and HHT in combination (n=10) for 25 days. Leukemic burden was monitored by peripheral blood (PB) draws and quantitation of leukemic cells (human CD45^+^, HLA-ABC^+^ cells) by flow cytometry on the indicated days is shown.

Finally, we wanted to determine if CB-839 plus HHT combination therapy can show efficacy in treating a different type of leukemia, acute lymphoblastic leukemia (ALL). Thus, we tested the effects of the drug combination using the Z-119 ALL cell line. Similar to what was observed in AML cells, HHT induced a greater amount of mitoROS when used in combination with CB-839 (Figure 7A), which correlated with decreased cell viability (Figure 7B) and increased apoptosis (Figure 7C). Comparable effects on cell viability (Figure S5A) and apoptosis (Figure S5B) were observed using a second ALL cell line, SUP-B15. To determine if CB-839 plus HHT shows efficacy in treating ALL *in vivo*, we utilized a mouse model of Bcr-Abl^+^ B-cell ALL as described previously^21^. Bl6 mice were transplanted with mouse ARF^−/−^ p185 Bcr-Abl^+^ B-ALL cells and after 3 days to allow for engraftment, mice began therapy with vehicle, HHT, CB-839, or HHT and CB-839 in combination (n=5 per group). After 7 days of therapy, leukemic cells in PB were quantified by flow cytometry (GFP^+^/B220^+^) [Note that 3 mice in both the vehicle and CB-839 alone groups had already succumbed to leukemia at this point]. Figure 7D shows that while CB-839 by itself had no significant effect, HHT in combination with CB-839 was far more effective than HHT alone in controlling leukemic burden (as shown on days 7 and 10), which resulted in significant extension of survival (Figure 7E). Taken all together, these results strongly argue that the strategy of combining a glutaminase inhibitor, such as CB-839, together with a pro-oxidant drug, such as HHT or ATO, has substantial therapeutic potential for treating multiple types of leukemia.

**Figure 7.**
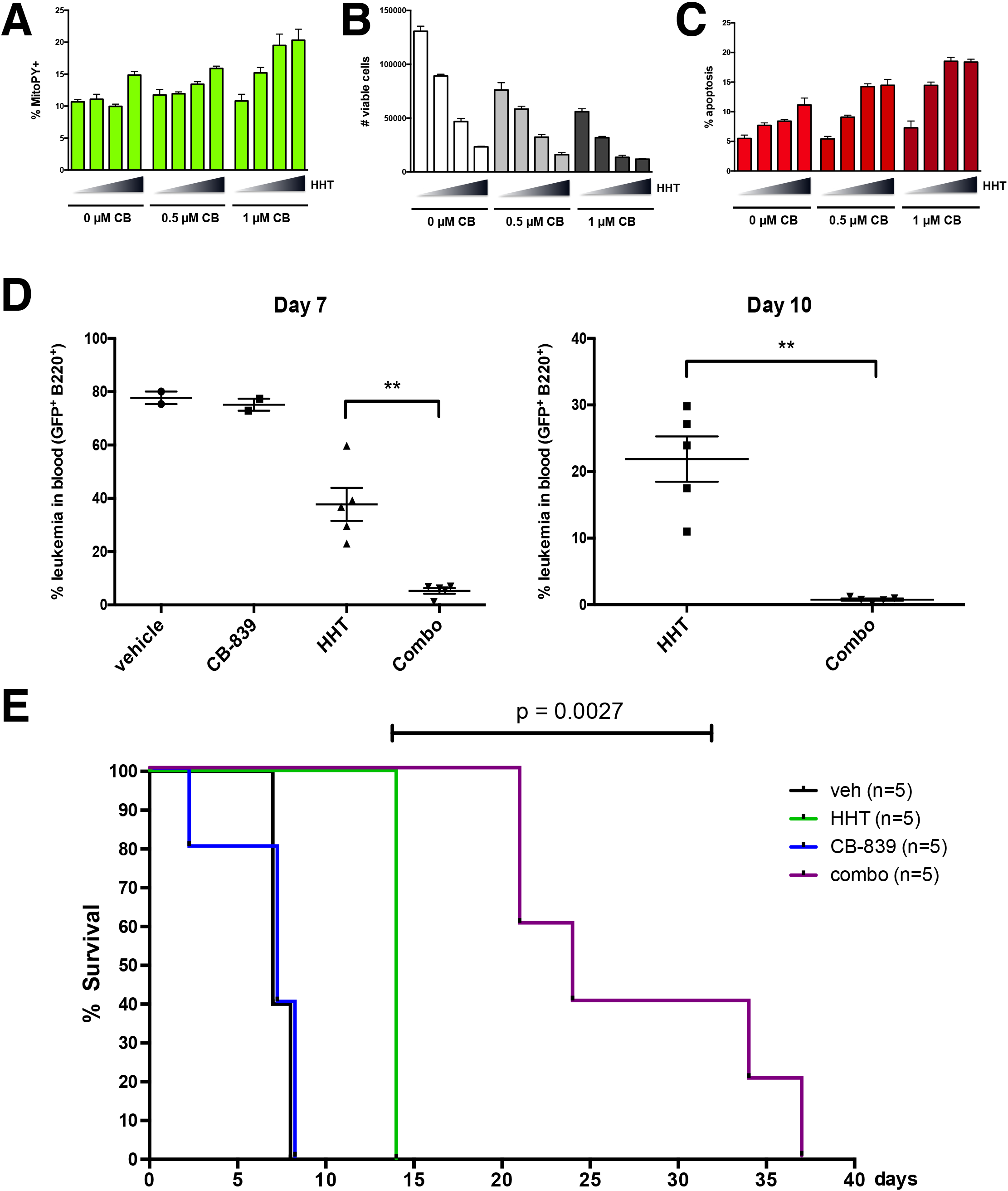
CB-839 cooperates with HHT in eliminating ALL cells *in* vitro and *in vivo*. **A, B, C)** Z-119 B-ALL cells were treated with CB-839 and HHT alone or together as indicated for 36 h (triangles: 0, 5, 10, and 20 nM HHT) and mitoROS levels were measured using MitoPY1 (**A**). After 72h, the number of viable cells (based on PI exclusion) was counted by flow cytometry (**B**) and apoptosis was measured after 48 h (**C**). **D)** ARF-/- p185 Bcr-Abl/GFP mouse B-ALL cells were transplanted into Bl6 mice and after 5 days, treatment was initiated with vehicle (n=5), CB-839 (n=5; 200 mg/kg twice daily p.o.), HHT (n=5; 1 mg/kg once daily i.p.), or CB-839 and HHT in combination (n=5) for up to 15 days (5 days on/2 days off therapy). After 7 and 10 days of therapy, peripheral blood from all mice was immunostained for B220 and Mac-1 and analyzed by flow cytometry. The percentage of GFP^+^ cells in the B-lineage (B220^+^, Mac-1^−^) population was determined and plotted. **E)** Kaplan-Meier curve showing survival of mice receiving the indicated therapy. Mice were sacrificed when moribund, and all showed clear evidence of leukemia in blood, bone marrow and spleen. Statistical significance determined using the log-rank (Mantel-Cox) test.

## Discussion

It has long been recognized that tumor cells exhibit metabolic characteristics that distinguish them from normal differentiated non-proliferative cells ^7–9^. The altered glucose metabolism that is frequently observed in tumors has been extensively studied and is already being exploited for cancer therapy ^25^ Altered glutamine metabolism in cancer, however, has only recently begun to be investigated. Glutamine is the most abundant amino acid in plasma and most tumors utilize glutamine at a much higher rate compared to normal cells. Glutamine metabolism contributes to energy production, macromolecular synthesis, and redox homeostasis, and is essential for survival of some cancer cells that have become addicted to glutamine ^10,11,26^.

Several recent studies have shown that targeting glutamine uptake or metabolism can have anti-leukemic activity in AML ^13,15,16,27,28^. However, most of these studies have focused on the role of glutamine in supporting energy metabolism rather on its role in maintaining redox homeostasis. Our recent studies of FLT3-mutated AML have shown that inhibition of FLT3 severely impairs glutamine metabolism, which leads to a deficiency in glutathione levels and accumulation of mitoROS that is causative in apoptotic cell death ^14^ Similar effects were observed in FLT3-mutated AML cells upon direct inhibition of glutaminase ^15^. In this report, we show that a glutaminase inhibitor causes decreased glutathione levels and increased mitoROS and apoptosis in an array of different AML cell lines, including those that are FLT3 WT. Furthermore, we show that addition of exogenous glutathione, which has no effect on energy production, is sufficient to suppress mitoROS levels and apoptosis upon glutaminase inhibition, strongly arguing that glutathione-dependent redox homeostasis is critical for maintaining cell survival in AML. Importantly, this dependence on glutamine/redox metabolism is independent of FLT3 status, and thus glutaminase inhibitors may have broad therapeutic utility in the treatment of various types of AML.

Numerous redox-directed therapeutic agents have been investigated in recent years for the treatment of cancer, so far with limited success, at least as monotherapy ^29–32^. Given that glutaminase inhibition leads to a compromised redox state, we surmised that this would sensitize the cells to adjuvant prooxidant drugs, leading to more apoptosis and more complete leukemia cell elimination. Indeed, we show that two different leukemia drugs that are potent inducers of mitoROS, ATO and HHT, have enhanced anti-leukemic activity in AML when combined with the glutaminase inhibitor CB-839, both *in vitro* and *in vivo*. Additionally, we show that CB-839 in combination with HHT shows efficacy in treating a different form of leukemia, ALL. Other hematological malignancies, such as multiple myeloma, have been reported to be highly dependent on glutamine metabolism^33 34^. Furthermore, subsets of many solid tumor types, including lung, breast, kidney, and prostate cancer, have been shown to be susceptible to glutaminase inhibition ^35–40^. Therefore, the use of a glutaminase inhibitor in combination with a pro-oxidant drug could represent a strategy with broad therapeutic utility in the treatment of cancer.

## Acknowledgements

We thank Dan Pollyea and Craig T. Jordan (University of Colorado SOM) for providing primary AML samples, and Craig Jordan for critical review of the manuscript. We thank Timothy Pardee (Wake Forest SOM) and Richard T. Williams (St. Jude Children’s Research Hospital) for the MLL-ENL/FLT3-ITD^+^ AML and Bcr-Abl^+^ ALL mouse models, respectively. Grants from the NIH/NCI (K22-CA133182) and the Cancer League of Colorado to M.A.G, and the V Foundation (T2016-012) to J.D., supported these studies. A.D. was supported by funds from the Boettcher Foundation, Webb-Waring Early Career award 2017.

## Authorship and conflict-of-interest statement

Conceptualization, M.A.G. and J.D.; Methodology, M.A.G., T.N., and A.D.; Data analysis, M.A.G., T.N., V.Z. and S.G.; Investigation, M.A.G., T.N., V.Z., and H.J.P.; Writing - Original Draft, M.A.G. and J.D.; Writing - Review & Editing, A.D.; Supervision, M.A.G., K.C.H., A.D., and J.D.; Funding acquisition, M.A.G., A.D., and J.D.

The authors declare no competing financial interests.

